## Supplementary materials for "A method to estimate the cellular composition of the mouse brain from heterogeneous datasets"

### Supplementary methods

#### Orientations field and depth processing

Processing the distance towards locations in the BBCA is crucial for *in-silico* modeling. For instance, assigning morphologies to cells requires knowing the space available for them [1]. Similarly, several literature reports depth-based density [2] or connectivity rules [3], which makes it more difficult to integrate them into brain models. In this paper, we validated our density profiles of the barrel field against the results from Meyer [4] which requires to compute the distance of each voxel of the AV towards the pia, guided by the orientation field of the main fibers axis in the isocortex (see S5 Fig. D).

To obtain this orientation field, we created a semi-automated method to generate coordinate systems using only a user-defined list of brain regions as reference. The goal was to compute a vector field pointing towards a general direction, based on a source region, a target region. Within the region of interest, direction vectors are obtained as the normalized gradient of a scalar field. Which in turn is obtained by assigning to every voxel of the brain a user-defined weight representing its distance from the source region (see S5 Fig. AB). For each subregion of interest, a single weight is assigned to every voxel of that subregion and a default value is given to the rest of the brain. For the isocortex, the following weights were assigned to the voxel of the scalar field:

- -2 for the lateral forebrain bundle system (corpus callosum)
- 1 for the layer 6
- 2 for the layer 5
- 3 for the layer 4
- 4 for the layer 3
- 5 for the layer 2 (and 2 / 3)
- 6 for the layer 1
- 0 for the other brain regions

To avoid boundary effects from the outside at the borders of a region, we extended the scalar field of the region to its surroundings voxels (see S5 Fig. B). These surrounding voxels are detected by a shading algorithm which looks for voxels close to annotation borders (i.e., where a change of annotation is occurring). We applied this shading algorithm to each layer of the isocortex, setting their surrounding voxels to the same weight in the scalar field.

An additional scalar shading is computed based on the distance to a subregion of interest identified as a target for fibers. This shading was created to attract the gradient of the voxels in the target region towards the outside. For the isocortex, voxels closed to the L1 and outside of the brain were assigned to 6 plus their distance to L1.

A Gaussian filter is then applied to the initialized scalar field and the gradient of the normalized blurred scalar field is eventually returned. The direction vectors are given by this gradient (see S5 Fig. C). This process was applied to all cortical areas by defining the white matter as the source region, and the outside of the brain as the target.

The orientation field algorithm can also be applied to the subregions of the cornu ammonis (CA), where the source and target regions are defined respectively as the stratum radiatum and stratum oriens. Similarly, it could be used in the Cerebellar Cortex defining the source region as the arbor vitae and the target region as the molecular layer.

The process of finding depth and boundaries for a voxel follows a straightforward procedure:

- Starting from the position of the voxel, add the unit vector corresponding to the orientation field axis (direction of pia) and check the layer at the new position.
- If the layer is different after the step, a boundary has been crossed and we record the distance traveled to get to it.
- Repeat this process until the isocortex has been exited.

The same can be done by subtracting the unit vector from the position (direction of white matter) to get the locations of the boundaries of deeper layers. Finally, a mean layer boundary distance to the pia can be extracted from the distance of its voxels.

#### Review of literature on densities of inhibitory neuron in the mouse brain

We performed a systematic review of the literature for inhibitory neurons in the mouse brain. To do so, we mostly leveraged the google scholar tool to find articles. Our searches involved combinations of the following key words (and their respective common abbreviations, e.g.: GAD67 for GABAergic): mouse, neurons, cells, inhibitory, GABAergic, parvalbumin, somatostatin, vasoactive intestinal peptide, interneurons, quantitative analysis, counts, densities, etc. We additionally searched more specifically for specific regions of the brain to confirm findings or try to improve our coverage of the whole brain. Except for the striatum, we only selected papers on mice experiments. We also made sure to select a single paper per experiment to prevent any duplicate referencing. We extracted the mean value provided in each paper and its standard deviation whenever this value was available, Standard Error of the Mean were converted to standard deviations based on the numbers of individuals used to make the estimates. A density estimate in a specific brain region was assumed to be valid for the entire region. If the estimates were provided in the form of counts, we converted these into densities using the region volume from the paper, if available, or computed from the AV. When ratios (or percentages) of neuron type in a region were provided as proportions of the local cells or neurons populations, these ratios were multiplied by the corresponding densities from the BBCAv1 [5]. In total, we extracted density values from 54 different papers [2,3,5–56]. Our literature review is compiling the work of many papers. Any use of the results presented in this review should give the credit to the corresponding paper these results were extracted from. In particular, the major contributor of this review is the Kim et al. [6] study, and it should get a particular attention when our work is cited.

For each paper providing regional counts, densities, or proportion of inhibitory neuron types, we assigned the closest matching AV region(s). Sometimes several regions were assigned to the same literature value, while sometimes multiple values from literature were averaged to fit into a region of the AV. For instance, the cortical layer 6 is subdivided into two sublayers a and b in the AV but some papers do not make this distinction [7,52]. For these cases, we assumed the same densities for both subregions.

Several papers provided densities for the frontal cortex [50–52]. This region was assumed to cover the prelimbic area, the frontal pole and infralimbic area of the AV.

Almási et al. [7] provided inhibitory neuron densities in the somatosensory cortex, barrel field. The estimates from the layer 5a and 5b in the paper were averaged as the AV does not have this layer subdivision.

Bjerke et al. [9] stored their PV+ neuron density values in their supplementary Table 1.

From Fasulo et al. [11], we obtained PV+ neuron densities from their Figure 1CE.

Fazzari et al.'s counts of inhibitory neurons in 40 μm coronal slices of the isocortex [12] were extracted from the paper's Figure 1.

We extracted PV+ neuron densities from the Figure 7 of Förster [13].

Gonchar et al. reported densities that we extracted from their Figure 2 [14].

In Gotts et al. [15], GAD67 immunoreactive neurons were counted in 50 μm sections of the nucleus of the solitary tract of the mouse brain (see their Table 1). We converted these counts into densities, using the mean volume of 50 μm coronal slices of the nucleus of the solitary tract in the AV. The paper's estimates in the subregions of the nucleus of the solitary tract were matched with their closest region in the AV:

- Medial part (paper) → Gelatinous part (AV)
- Central part → Central part
- Commissural part → Commissural part
- Dorsomedial, Intermediate, Medial parts → Medial part
- Interstitial, Lateral, Ventrolateral, Ventral parts → Lateral part

We took the sum of the left and right hemisphere as this distinction does not exist in the AV. We also averaged all values assigned to the lateral and medial subregions in the AV.

The globus pallidus densities in Gourfinkel-An et al. [16] were assumed to be the same in both its internal and external segment subdivisions in the AV. A global estimate of inhibitory neurons in the entire cerebellar cortex was estimated from Gourfinkel-An et al. based on the authors estimates in the molecular and granular layers. These estimates were multiplied by the proportion of each layer volume in the cerebellar cortex. No voxels of the AV are labelled as belonging to the Purkinje Layer so its contribution to the cerebellar cortex density estimate was null.

Grünewald et al. [17] supplementary Table 2 lists all the interneuron densities reported by the authors.

Hafner et al. [18] provided counts of VIP+ neurons in the somatosensory cortex, barrel field. These counts were converted into densities using the size of the area used to count the cells (0.1 mm * 0.24 mm) multiplied by the mean depth of the region, based on measurements from Lefort et al. [24].

Han et al. [19] provided counts of GAD67 neurons from whole brain 30 µm-thick coronal slices. The mean surface occupied by the regions of the article in the coronal slices were estimated from similar coronal slices in the AV. The resulting densities are very low when compared to other literature sources. These values were therefore not considered in the fitting process.

Irintchev et al. [21] reported inhibitory neuron densities which we extracted from their Fig. 3. However, it was not very clear from which region of the somatosensory cortex one of their estimates were taken from. Hence, we considered these estimates for the entire region.

Jinno and Kosaka [22] and Whissell et al. [51] reported densities in the dorsal and ventral parts of the hippocampus. This subdivision of the hippocampus is not present in the AV, and as a result, the values reports were averaged to represent the full region. Additionally, estimates from the lateral and medial subdivisions of the visual areas from Whissell et al. were averaged for similar reasons.

Lefort et al. [24] provided counts of inhibitory neurons in a cylinder of the C2 barrel column of the mouse isocortex. We transformed the counts into densities based on the dimensions of the cylinder provided by the authors. We additionally applied the densities in the C2 barrel column to the whole barrel region (including the septa).

Moreno-Gonzalez et al. [26] provided densities of neurons in the Entorhinal area in the paper Figure 1.

We extracted manually the densities from the Figure 11 in Neddens and Buonanno [27].

In Okada et al. [29], densities of GAD67 neurons in the mouse nucleus of the solitary tract were measured by the authors in 4 areas of the regions. The average of the densities in the areas was taken, as the areas taken by the authors did not correspond to any of the AV subregions.

Ono et al. [30] provided densities of GABAergic neurons in cells / $10^4$ $\mu m^2$ from 50 μm-thick transverse slices of the mouse brain. No standard deviation value was provided in the paper.

Parrish-Aungust et al. [31] obtained multiple counts of inhibitory cells in their Table 4 that were converted into densities using the volumes of the regions provided by the authors in Table 2.

In Pirone et al. [32], the authors reported densities of inhibitory neurons in coronal sections from the Infralimbic and Prelimbic areas. We converted these densities in cells/$mm^3$ based on the thickness of the sections (20 μm).

Pitts et al. [33] reported densities of cells/$mm^2$ in 40 μm-thick slices in their Figure 3B.

Prönneke et al. [34] have reports of VIP neuron densities in the somatosensory cortex, barrel field.

From Ramos et al. [36], we extracted densities of SST+ neurons in different regions of the hippocampus from their figure 2E.

In the Figure 8, of Ransome and Turnley [37], we could obtain densities from the Striatum.

Sanchez-Meijas et al. [38] reported densities of PV+ neurons from two regions of the cortex. We considered the Zone 35, and Zone 36 (in the paper) as respectively perirhinal area and the ectorhinal area of the isocortex.

Schmalbach et al. [39] have neuron densities reported that we extracted from their Figure 1 and 2.

Schmid et al [40] provided densities in cell/$mm^2$ from 50 μm-thick slices in their Figure 1i.

Song et al. [42] provided counts of PV+ neurons (see their table 1) in the striatum together with the volume in which the counting was performed.

We extracted density values from the Table 2 of Suzuki and Bekkers [43] in the anterior part of the piriform area of the olfactory areas (i.e. the part in contact with the lateral olfactory tracts).

In absence of a better We assumed these estimates to be true in the entire piriform area, The layer 1, 2 and 3 were respectively associated to the molecular layer, the pyramidal layer and the polymorph layer. The a and b subdivisions of the layers 1 and 2 were not present in the AV so we averaged the two estimates for each layer.

Tamamaki et al. provided percentages of GABAergic cells among the population of neurons in regions of the isocortex [44]. We converted these proportions into densities using the neuron densities from the Cell Atlas.

A proportion of GABAergic neuron densities in the striatum were estimated from a collection of papers on the rodent [57–62], see review from Tepper et al. [45]. From these papers, we could estimate the total proportion of inhibitory neuron in the striatum as the sum of the proportions of each of its distinct GABAergic cell type:

- Medium spiny cells: 95% [57].
- Cholinergic: 1.7% [58].
- Parvalbumin: 0.7% [59,60].
- Calretinin: 0.8% [59].
- Neuropeptide Y: 0.9% [61].
- Tyrosine hydroxylase: 0.2% [62].

The total percent of inhibitory neuron in the striatum is therefore at least equal to 99.3%, which is the value we used in our pipeline.

We extracted PV and SST neuron densities from the main text and the Figure 1 of the paper of Trujilo-Estrada et al. [46].

We extracted from Figure 3 and 4 of Waider et al. [47] densities of PV expressing neurons.

Counts of GAD67 neurons in the mouse nucleus of the solitary tract from 10 sections (100 * 100 * 50 µm) were manually extracted from the Fig. 1F in Wang and Bradley [49].

Wang et al. [50] reported percent of inhibitory neuron cells according to the total cell population in their Figure 4. We used the BBCAV1 cell densities to convert this proportion into densities.

Xu et al. [52] reported densities in the layer 4 of the frontal cortex. But this layer does not appear in the frontal part of the cortex of the AV, which is why the value was ignored.

We extracted counts of PV+ neurons in the striatum from Yalcin-Cakmakli et al. [53]. To convert these counts into densities we used the volume of the striatum from the AV2a.

From the paper of Zhang et al. [55] we extracted the density values present in their Figure 6.

From Zhao et al. [56] Fig. 3 we manually extracted the counts and standard deviation of GAD67 positive cells in coronal slices of the lateral septal nucleus. The subregions from the paper are more precise than the AV. We averaged the intermediate counts values with the rostro-ventral counts and assigned them to the rostroventral region of the AV. The caudal-ventral counts from the paper were assigned to the ventral part of the AV as the voxels assigned to this region are located at the most caudal part of the lateral septal nucleus. The values left of the paper were averaged and assigned to the caudal-dorsal part of the AV. Counts were then converted to densities based on the volume of the counting frame (i.e., 163 * 163 * 30 µm).

From Erö et al. [5], we finally extracted the neuron densities in regions reported in literature to be fully inhibitory. These includes the layer 1 of the isocortex [35], the molecular layer of the cerebellar cortex [3], the reticular nucleus of the thalamus [20].

#### Finding an initial solution for the optimization

After obtaining first estimates of counts, for each inhibitory neuron type, in each region of the brain, from the literature or the fitting, we need to test and correct them using the assumptions defined in the Assumptions section. There are two consequences of assumptions 2 and 3: First, the number of GAD67 positive neurons in each brain region must be smaller than the total number of neurons (computed at step 2 of the BBCAv2 pipeline - see Fig 1) and second, the sum of VIP, SST and PV positive neurons must also be smaller than the total number of neurons. Thus, whenever these conditions are violated, we scale down the number of these neurons while preserving their ratios (see Algorithm 1: S2 Fig). This correction is applied in each region starting from leaf regions in the AV hierarchy to the top-level brain regions. Leaf regions are independent so corrections can be applied directly. For regions higher up in the hierarchy (e.g., barrel cortex), the estimated cell counts must equal the sum of the cell counts of their subregions (e.g., layers of barrel cortex). Solving this constraint is therefore easier following this order.

Then, we assert that our inhibitory neuron density estimates are taking into account our 4th assumption which states that PV, SST and VIP also express GAD67 and allows us to estimate nRest. Thus, for each region of the brain following the same order as described previously, whenever the constraint is violated, we scale down the number of the PV+ SST+ and VIP+ neurons while preserving their relative ratios and conversely increase the number of GAD67+ neurons (see Algorithm 2: S3 Fig).

Algorithms 1 and 2 are applied to each region of the AV starting from leaf regions in the hierarchy tree to the major brain regions. This implies that the corrections applied to a region are impacted by the ones applied on its subregions but not the other way around. Hence, these algorithms are appropriate to find a good initial solution for the Cell Atlas model (see combination section) but does not guarantee an optimal result, i.e., as closed as possible from the literature and the fitting original estimates.

#### Supplementary figures

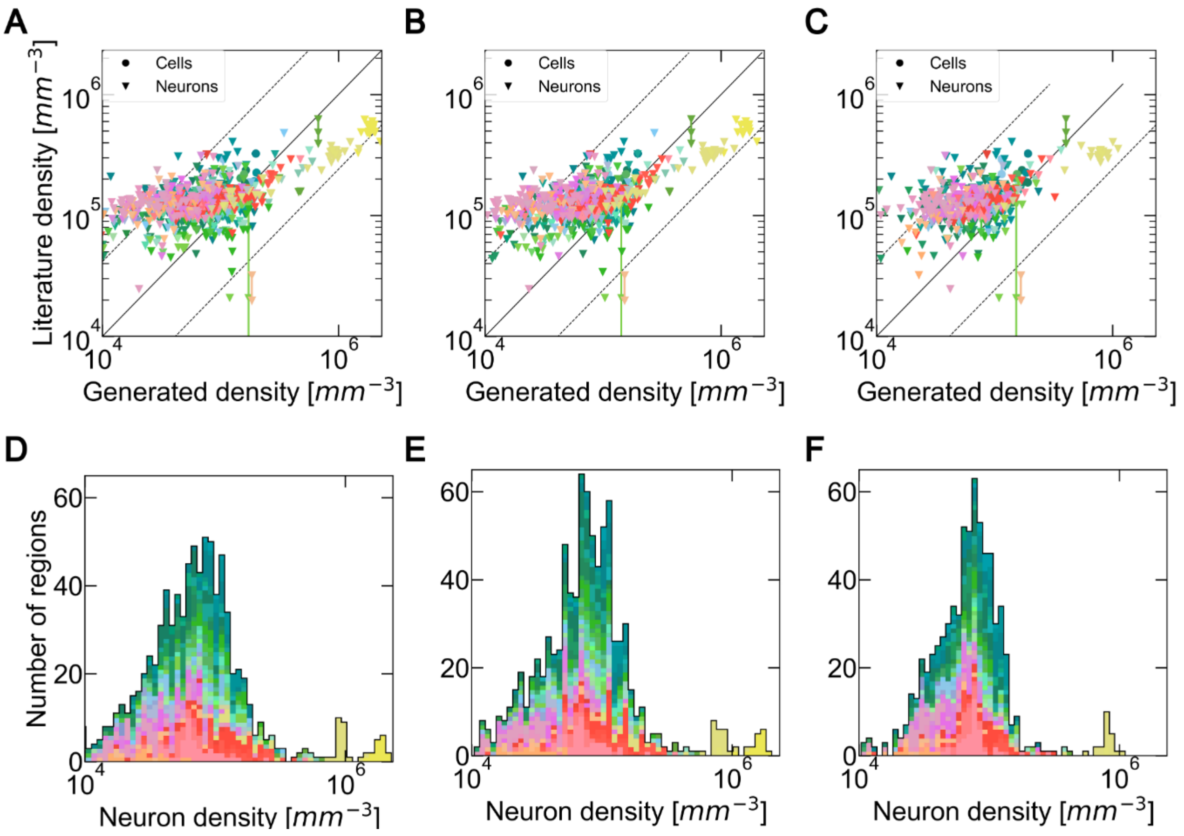


**S1 Fig. Global impact of the annotation atlas on cells and neurons densities**

(A), (B), (C) Distribution of cell and neuron density values reported in literature against generated densities for similar regions, using the 3 pairs of annotations and Nissl volumes (from left to right: CCFbbp, CCFv2 from the AIBS, CCFv3 from the AIBS). When multiple literature sources are available for the exact same region, they are both shown as a data point and are linked together. The color encodes the brain regions according to the AV, while the shapes of the points encode for cell types. The middle line delimits equal quantities, while the dashed line shows the average deviation of 2.7-fold between literature values reporting on the same region. Some subregions of the brain are not represented in CCFv3 which explains the different numbers of points.

(D), (E), (F) Histogram of the brain regions in terms of neuron density values for the 3 pairs of AV and Nissl volumes (from left to right: CCFbbp, CCFv2 from the AIBS, CCFv3 from the AIBS). Each region is represented with a single-color patch of the same size.

We ordered $\mathbf{R}$ according to the regions depth in the region hierarchy of the AV to obtain $\boldsymbol{R_O}$.
Confidence interval of $nPV$ (similarly for $nSST$, $nVIP$, and $nGAD$), $\forall r \in \mathbf{R}$:

$$[minPV_r, maxPV_r] \text{ where } \begin{cases} minPV_r = nPV_r - stdPV_r \\ maxPV_r = nPV_r + stdPV_r \end{cases}$$

```
foreach r ∈ R_O do
    if nNeu_r < nGAD_r then
        nGAD_r = nNeu_r
    end
    maxGAD_r = min(maxGAD_r, nNeu_r)
    minGAD_r = max(minGAD_r, nNeu_r)
    sumPSV = nPV_r + nSST_r + nVIP_r
    if nNeu_r < minPV_r + minSST_r + minVIP_r then
        /* case where no solution can be found within confidence intervals,
           PV, SST and VIP proportions are kept                              */
        nPV_r =  minPV_r =  maxPV_r = nNeu_r × nPV_r / sumPSV_r
        nSST_r = minSST_r = maxSST_r = nNeu_r × nSST_r / sumPSV_r
        nVIP_r = minVIP_r = maxVIP_r = nNeu_r × nVIP_r / sumPSV_r
    end
    else if  nNeu_r < sumPSV then
        /* case where there is a solution within confidence intervals,
           but mean estimates are greater than the number of neuron          */
        remainingSum = sumPSV
        while nNeu_r) < sumPSV do
            p_PV = nPV_r / remainingSum
            p_SST = nSST_r / remainingSum
            p_VIP = nVIP_r / remainingSum
            diff = sumPSV − nNeu_r
            nPV_r = max(minPV_r,  nPV_r − p_PV × diff)
            nSST_r = max(minSST_r,  nSST_r − p_SST × diff)
            nVIP_r = max(minVIP_r,  nVIP_r − p_VIP × diff)
            remainingSum = sumPSV = nPV_r + nSST_r + nVIP_r
            if nPV_r = minPV_r then
                remainingSum = remainingSum − nPV_r
            end
            if nSST_r = minSST_r then
                remainingSum = remainingSum − nSST_r
            end
            if nVIP_r = minVIP_r then
                remainingSum = remainingSum − nVIP_r
            end
        end
        maxPV_r = nPV_r
        maxSST_r = nSST_r
        maxVIP_r = nVIP_r
    end
    else
        /* case where the mean estimates are correct,
           only maximums have to be checked.                                 */
        maxPV_r = min(maxPV_r, nNeu_r − nSST_r − nVIP_r)
        maxSST_r = min(maxSST_r, nNeu_r − nPV_r − nVIP_r)
        maxVIP_r = min(maxVIP_r, nNeu_r − nPV_r − nSST_r)
end
```

#### S2 Fig. Algorithm 1: Cap inhibitory densities to number of neurons

The estimated counts of GAD67+ neurons and the sum of PV+, SST+, VIP+ neurons counts are limited by the previously computed neuron counts (step 2 of the BBCAv2 pipeline - see Fig. 1) to ensure that assumptions 2 and 3 are fulfilled (see Section 2.4). Recall that **R** is the set of brain regions, ∀ r ∈ **R** inversely ordered according to their depth in the region hierarchy of the AV ($\mathbf{R_o}$), the algorithm checks if the conditions $nNeu_r \geq nGAD_r$ and $nNeu_r \geq nPV_r$

+ $nSST_r$ + $nVIP_r$ are satisfied. If not, it finds a solution which tries to match the following properties, ordered by priority: (1): Remain in range of the standard deviation of each value (confidence intervals), (2): Maintain the proportion of PV, SST and VIP within the region. If a solution exists within the confidence intervals, then the number of extra neurons (diff variable at line 19) is subtracted proportionally to the ratio of each neuron type (computed at lines 16-18). Then, if one of the neuron type estimates reaches the minimum of its confidence interval, then it is no more reduced (lines 20-22) and the remaining extra neurons are subtracted from the other neuron type estimates (lines 25, 28, 31).

We ordered **R** according to the regions depth in the region hierarchy of the AV to obtain $\boldsymbol{R_O}$.
Confidence interval of $nPV$ (similarly for $nSST$, $nVIP$, and $nGAD$), $\forall r \in \mathbf{R}$:

$$[minPV_r, maxPV_r] \text{ where } \begin{cases} minPV_r = nPV_r - stdPV_r \\ maxPV_r = nPV_r + stdPV_r \end{cases}$$

**foreach** $r \in \boldsymbol{R_O}$ **do**
    $sumPSV_r = nPV_r + nSST_r + nVIP_r$
    **if** $sumPSV_r > nGAD_r$ **then**
        $q = (sumPSV_r - nGAD_r) \;/\; (sumPSV_r - nGAD_r + maxGAD_r - minSumPSV_r)$
        // If $q \leq 1$ there is a solution within confidence intervals
        $nGAD_r = nGAD_r + q \times stdGAD_r$
        $nPV_r = nGAD_r \times nPV_r \;/\; sumPSV_r$
        $nSST_r = nGAD_r \times nSST_r \;/\; sumPSV_r$
        $nVIP_r = nGAD_r \times nVIP_r \;/\; sumPSV_r$
    **end**
    $maxGAD_r = min(maxGAD_r, nGAD_r)$
    $minGAD_r = max(minGAD_r, nGAD_r)$
    $maxPV_r = min(maxPV_r, nPV_r)$
    $minPV_r = max(minPV_r, nPV_r)$
    $maxSST_r = min(maxSST_r, nSST_r)$
    $minSST_r = max(minSST_r, nSST_r)$
    $maxVIP_r = min(maxVIP_r, nVIP_r)$
    $minVIP_r = max(minVIP_r, nVIP_r)$
**end**

#### S3 Fig. Algorithm 2: Maintain inhibitory densities coherence

This algorithm corrects the estimated densities of PV+, SST+, VIP+ and GAD67+ neurons in the model so that assumption 4 of Section 2.4 is fulfilled. **R** is the set of brain regions, ∀ r ∈ **R** inversely ordered according to their depth in the region hierarchy of the AV (**R**$_o$), the algorithm checks if the condition nGADr ≥ $nPV_r$ + $nSST_r$ + $nVIP_r$ is satisfied. If not, it finds a solution which tries to match the following properties, ordered by priority: (1): Remain in range of the standard deviation of each value (confidence intervals), (2): Maintain the proportion (ratios) of PV, SST and VIP. When a set of value exists so that the property (1) is fulfilled, there is a correction factor q ∈ [0,1] which corresponds to the fraction of standard deviation needed to guarantee that the sum of the inhibitory subtypes remains under the estimated count of inhibitory neurons and that each value remains within its confidence interval.

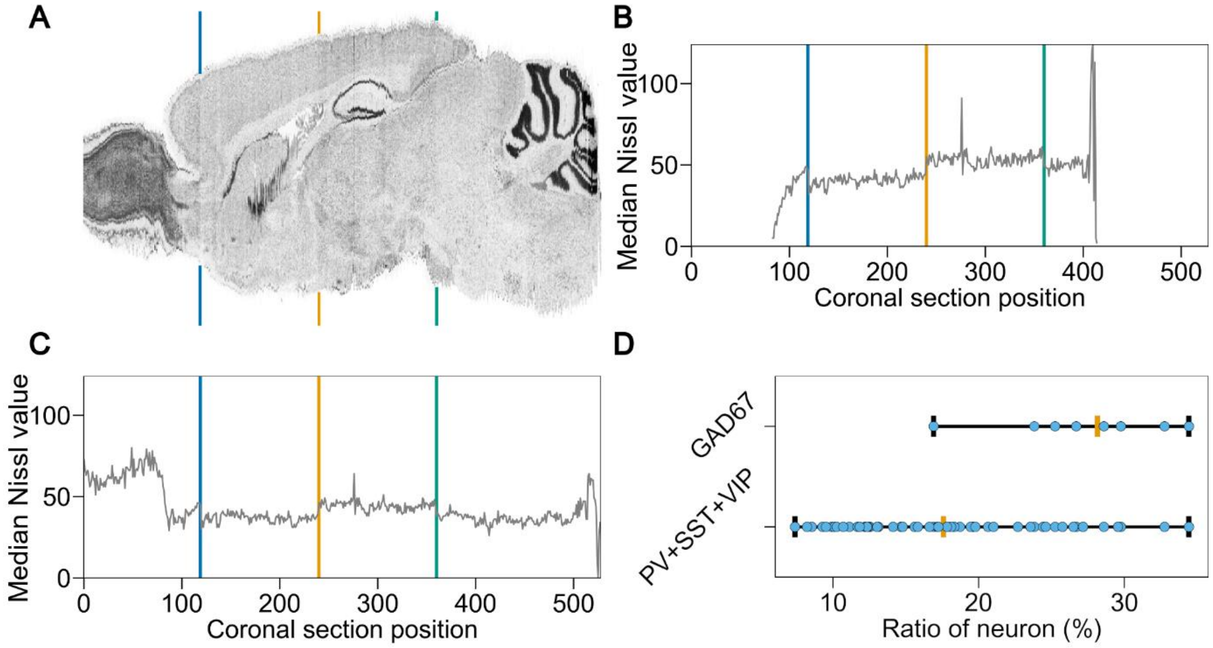


**S4 Fig. Explaining the isocortex inhibitory densities from BBCAv2.**

(A) Coronal view of the Nissl reference atlas used for BBCAv2, showing cells of the mouse brain. Regions with high cell density appear in dark grey. The blue, orange and green lines behind the sagittal slice highlight the rapid changes of Nissl expression coming from the original Nissl experiment from Dong [63]. (B) (C) Evolution of the median expression level along the sagittal axis, in the Nissl volume realigned in Erö et al. [5], for the whole brain (B) and the isocortex (C). The rapid changes of Nissl expression that were detected in (A) are also visible in (B) and (C).

(D) isocortex ratio of inhibitory neurons according to literature. The ranges show to ratios literature cell type counts in isocortex divided by the counts of neurons of the BBCAv2 in their corresponding region. The bottom and top distributions correspond to the proportion of respectively the sum of the reported values of PV, SST and VIP counts, and the reported values of GAD67. The mean value of each distribution is shown in orange. The minimum and maximum values are indicated by the whiskers.

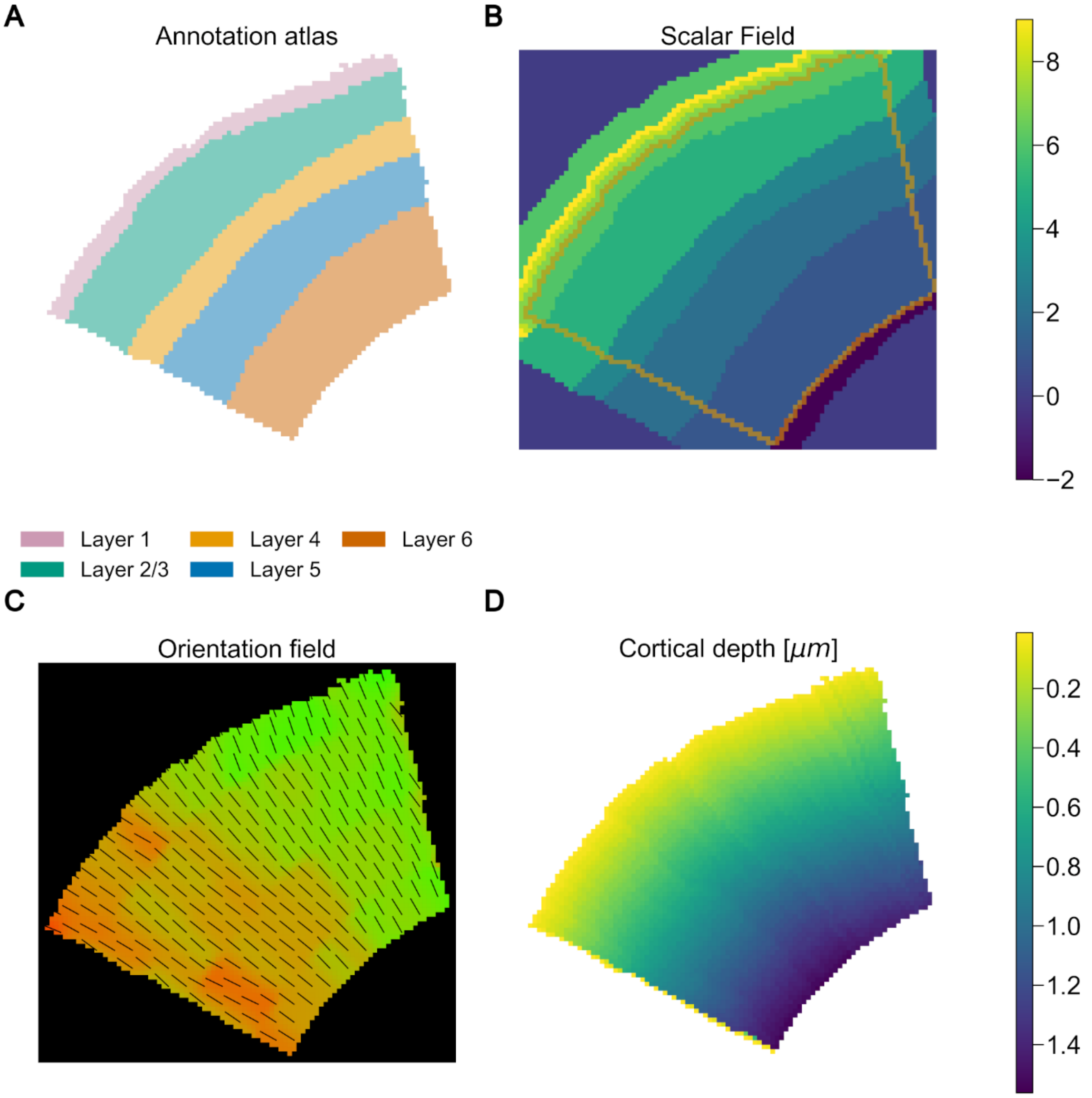


**S5 Fig. Orientations field and depth computation of the barrel cortex**

(A) Coronal slice of the AV showing the barrel cortex and its different sublayers.

(B) Coronal slice of the scalar field of the barrel cortex. A weight is assigned to every voxel of the AV. These weights follow the order of crossing the isocortex by its fibers from the corpus callosum to layer 1. The borders of the barrel field are highlighted in orange. The weight assigned to the surrounding voxels of the region corresponds to their closest layer's. The weight of the voxels outside the region beyond layer 1 increases as moving away from the barrel cortex.

(C) Coronal slice of the orientation field of the barrel cortex. To each voxel of the AV, a 3D direction normalized vector is computed corresponding to the main axis of the axons in the region. Colors represent the orientation vectors norm on their respective plane, black lines their projected axis.

(D) Coronal slice of the depth according to pia in the barrel cortex expressed in micrometers.

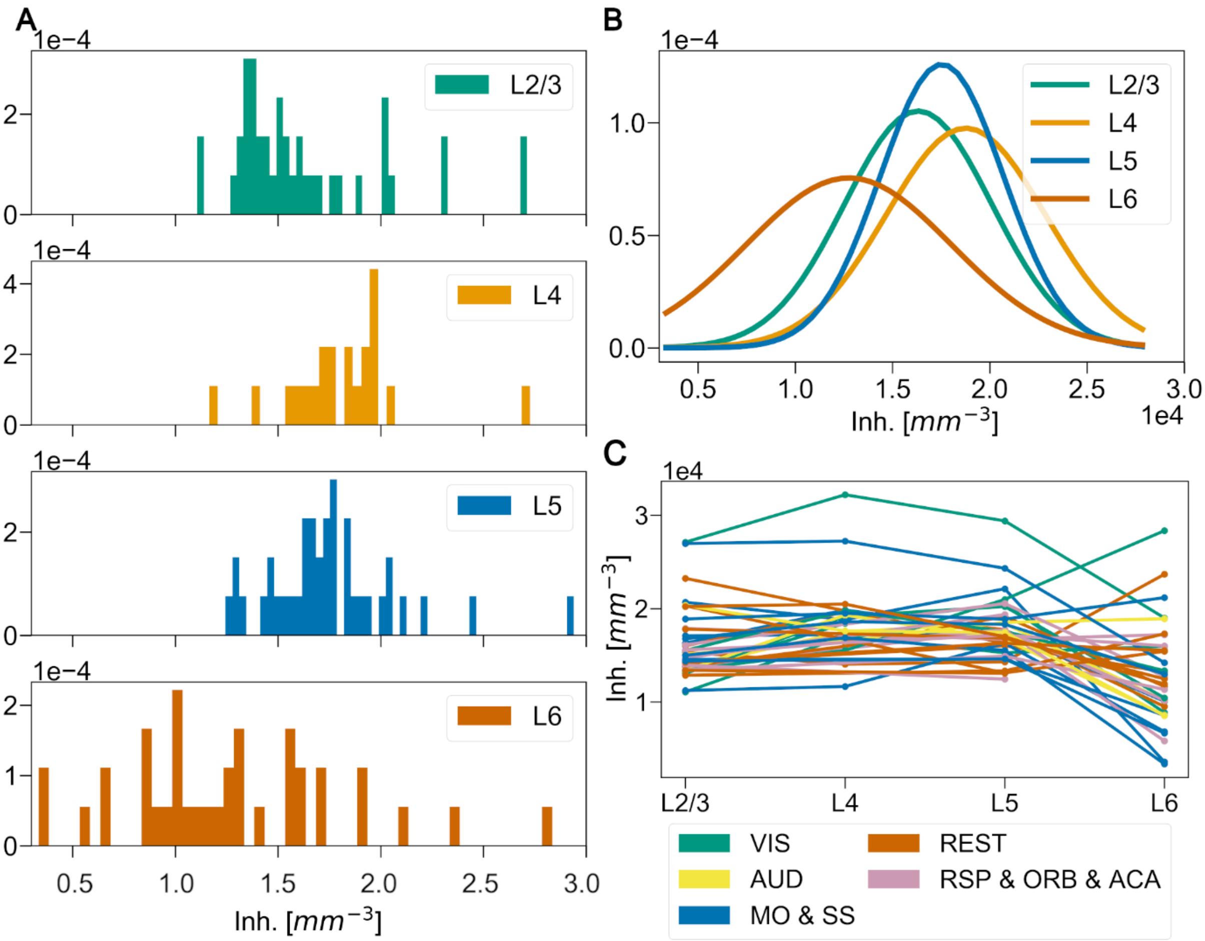


**S6 Fig. Cortical inhibitory density distribution**

(A) Normalized distribution of cortical inhibitory neuron densities across the isocortex subregions grouped by layers. For each layer, the fitted normal distribution associated is displayed as a line on top of it. The normal distributions of L2/3 (mean 1.6x104, std. 3.8 x103), L4 (mean 1.9x104, std. 4.1 x103) and L5 (mean 1.8x104, std. 3.1 x103) are overlapping significantly.

(B) Distribution of inhibitory neuron densities according to cortical layers for each subregion of the isocortex. For most of the cortical subregions, the density is constant from L2 to L5 and drops for L6.

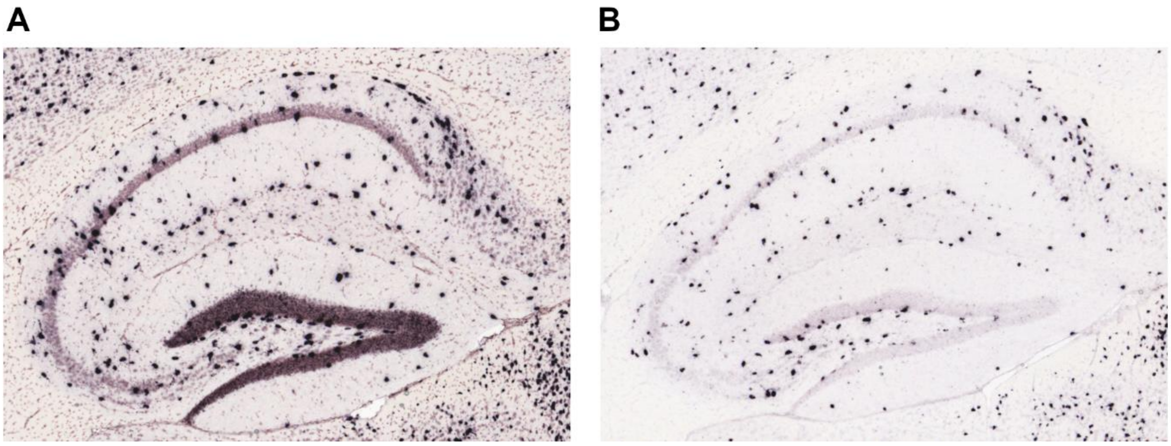


**S7 Fig. GAD67 and GAD65 ISH coronal slices showing the dentate gyrus**

GAD67 (A) and GAD65 (B) ISH coronal slices of the mouse brain from the AIBS website, showing the dentate gyrus (region resembling a greater-than symbol). Somas reacting to the marker stand out from the background. The more a cell is reacting to the marker the darker it will be shown in the image. Two different populations of cells reacting to the markers can be seen in the images. First, a population of cells with large somas, and strongly reacting to both the GAD67 and GAD65 markers. Second, a dense population of cells with small somas, reacting to the GAD67 marker but not to GAD65.

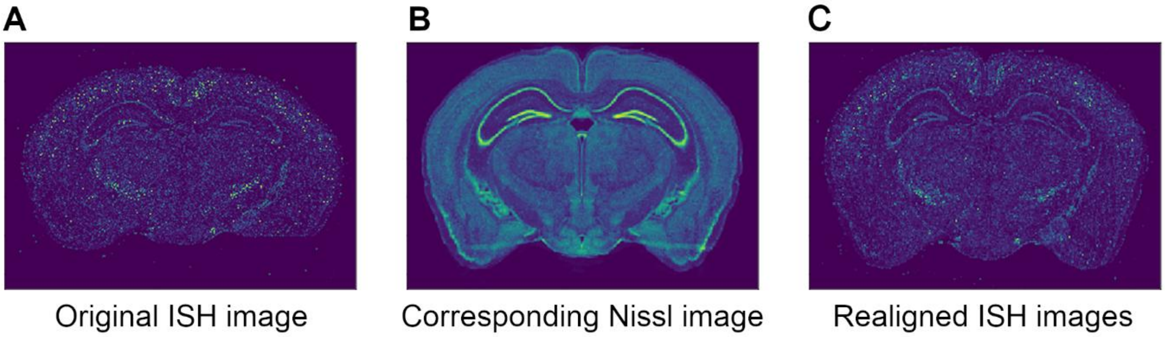


**S8 Fig. Results of the realignment of a PV ISH coronal slice to the Nissl volume**

This figure shows coronal slices of the mouse brain from AIBS experiments (Nissl and PV). ISH images from the AIBS (A) are realigned to their corresponding slice in the Nissl volume (B) using the Krepl et al.'s algorithm [64]. The registration is performed on the raw images from the AIBS as they provide more landmarks and then applied to the filtered images (C).

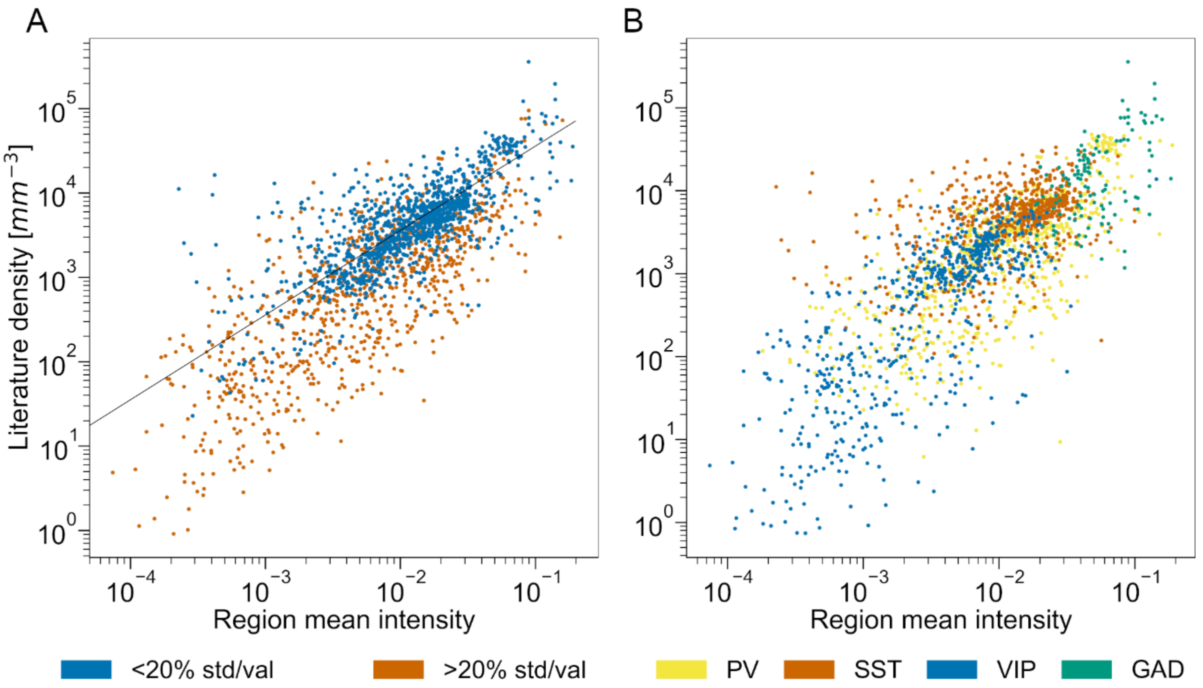


**S9 Fig. Linear fitting of marker intensity to cell density.**

Scatter plots of the PV+, SST+, VIP+ and GAD+ densities reported in literature (y-axis) according to the region mean intensity (x-axis). Each point represents a single literature density value. (A) The scatter plot is color-coded according to the different levels of confidence from literature data (ratio of standard deviation over mean value). The linear fit is represented with a black line. (B) Same scatter plot as (A) but the points are here color-coded according to the genetic marker expressed by the neuron population.

### Supplementary Tables

| Abbreviation | Full name |
|---|---|
| AAV | Adeno-associated virus |
| ACA | Anterior Cingulate Area |
| ACAd | Anterior cingulate area, dorsal part |
| ACAv | Anterior cingulate area, ventral part |
| AIBS | Allen Institute for Brain Science |
| AId | Agranular insular area, dorsal part |
| AIp | Agranular insular area, posterior part |
| AIv | Agranular insular area, ventral part |
| AUDd | Dorsal auditory area |
| AUDp | primary auditory cortex |
| AUDpo | Posterior auditory area |
| AUDv | Ventral auditory area |
| AV | Annotation Atlas Volume |
| BBCA | Blue Brain Mouse Cell Atlas pipeline |
| BF / SSp-bfd | Primary somatosensory area, barrel field |
| CA1 | Field CA1 |
| CA3 | Field CA3 |
| CBXpu | CereBellar corteX, Purkinje layer |
| CCF | Common Coordinate Framework |
| DG | Dentate Gyrus |
| ECT | Ectorhinal area |
| EXC | excitatory neurons |
| FRP | Frontal pole, cerebral cortex |
| GABA | gamma-Aminobutyric acid |
| GAD | glutamic acid decarboxylase |
| GU | Gustatory areas |
| ILA | Infralimbic area |

| | |
|---|---|
| INH | inhibitory neuron |
| ISH | *in situ* hybridization |
| LAMP5 | lysosomal associated membrane protein family member 5 |
| MOp | Primary motor area |
| MOs | Secondary motor area |
| ORBl | Orbital area, lateral part |
| ORBm | Orbital area, medial part |
| ORBvl | Orbital area, ventrolateral part |
| PERI | Perirhinal area |
| PL | PreLimbic area |
| PTLp | Posterior parietal association areas |
| PV | parvalbumin |
| RSPagl | Retrosplenial area, lateral agranular part |
| RSPd | Retrosplenial area, dorsal part |
| RSPv | Retrosplenial area, ventral part |
| SS / SSp | Primary SomatoSensory area |
| SSp-ll | primary somatosensory cortex lower limb |
| SSp-m | Primary somatosensory area, mouth |
| SSp-n | Primary somatosensory area, nose |
| SSp-tr | Primary somatosensory area, trunk |
| SSs | Supplemental somatosensory area |
| SST | somatostatin |
| STR | STRiatum |
| SUB | SUBiculum |
| TEa | temporal association areas |
| VIP | vasoactive intestinal peptide |
| VISal | Anterolateral visual area |
| VISam | Anteromedial visual area |
| VISC | Visceral area |
| VISl | Lateral visual area |

| | |
|---|---|
| VISp / VA | Primary VISual area |
| VISpl | Posterolateral visual area |
| VISpm | Posteromedial visual area |

**S1 Table. Nonstandard abbreviations**

List of all abbreviations used in this study. Region abbreviations are related to Fig. 4 and Fig. 8.

13. Förster JA. Quantitative morphological analysis of the neostriatum and the cerebellum of tenascin-C deficient mice. Quantitative morphologische Analysen des Neostriatums und

des Cerebellums der Tenascin-C defizienten Maus. 2008 [cited 18 Jan 2021]. Available: https://ediss.sub.uni-hamburg.de/handle/ediss/2354

38. Sanchez-Mejias E, Nuñez-Diaz C, Sanchez-Varo R, Gomez-Arboledas A, Garcia-Leon JA, Fernandez-Valenzuela JJ, et al. Distinct disease-sensitive GABAergic neurons in the

perirhinal cortex of Alzheimer's mice and patients. Brain Pathol. 2020;30: 345–363. doi:10.1111/bpa.12785

63. Dong HW. Allen reference atlas: a digital color brain atlas of the C57Black/6J male mouse. Hoboken, N.J.: Wiley; 2007.

64. Krepl J, Casalegno F, Delattre E, Erö C, Lu H, Keller D, et al. Supervised Learning With Perceptual Similarity for Multimodal Gene Expression Registration of a Mouse Brain Atlas. Front Neuroinformatics. 2021;15: 691918. doi:10.3389/fninf.2021.691918
